# A functional annotation based integration of different similarity measures for gene expressions

**DOI:** 10.64898/2026.02.23.707392

**Authors:** Sampa Misra, Sukriti Roy, Shubhra Sankar Ray

## Abstract

Genes with similar expression profiles often exhibit similar functional properties. An “integrated similarity score” (ISS) is developed by combining different expression similarity measures through weights, obtained using biological information, for improving gene similarity. The expression similarity measures are converted to the common framework of positive predictive value using functional annotation. A fitness function, called “fitness function using functional annotation of genes” (FFFAG), is also developed by minimizing the difference between functional similarity value and the ISS. The FFFAG is used to determine the weight combination of different similarity measures in ISS. In addition, an existing similarity measure, called TMJ (integrated similarity measure by multiplying Triangle and Jaccard similarity), is also modified to incorporate biological knowledge involving functional annotation. The results demonstrate that ISS is superior to individual similarity measure to find similar gene pairs. Further, the ISS predicts the functional categories of 40 unclassified yeast genes at *p*-value cutoff of 10^−10^ from 12 clusters. The associated code is accessible at http://www.isical.ac.in/∼shubhra/ISS.html.

## 1. Introduction

Microarray technology is one of the most powerful and commonly used techniques to monitor and analyze large quantities of genomic data efficiently and simultaneously [1; 2; 3]. In general, various types of analysis have been performed using gene expression data where identifying the group of genes with similar expression patterns is the key [1; 4]. Different methods of statistics, machine learning, and data mining have been used for gene expression analysis [5; 6]. The most useful unsupervised method is clustering to generate new hypothesis from gene expression data [7; 8]. In clustering problems, appropriate similarity measures are likely to produce clusters with higher number of functionally related genes as compared to other measures. These functionally related genes can be helpful in gene function prediction [9; 10; 11], functional module prediction [12; 13], gene regulatory network analysis [14], and disease prediction [15]. Thus, it is important to develop a similarity measure that can be helpful in finding functionally linked genes with similar gene expression values in an improved manner. Currently, the similarity between two genes is computed using Pearson correlation (PC) [16], Manhattan distance/Cityblock distance (MD) [3], Euclidean distance (ED) [17], and Spearman rank correlation (SRC) [18]. Boriah et al. [19] compared the similarity measures based on categorical data and evaluated them for outliers detection. Fernando et al. [20] compared and bench-marked binary-based similarity measures for categorical data. Deshpande et al. [21] used biological data to conduct a comparison for genetic interaction networks using profile similarity measures. They observed that the dot product is stable among different conditions for genetic interaction datasets.

ED mainly reflects the magnitude of all the changes in gene expression data without considering the shape [3]. PC is overly sensitive to the shape of gene expression data [3]. SRC [18] reflects the shape of the gene expression data. This rank-based method could mistakenly interpret a pattern as rank causes information loss. Costa et al. [22] compared three different distance measures for clustering of short time-series gene expression data. Gibbons and Roth [23] compared five different distance measures for gene time-series clustering. Although the methods developed in [22; 23] are used for clustering gene expression data, however, it is not clear from these investigations the most appropriate one. Since, these measures can not reflect the magnitude similarity at various time points and the shape similarity of expression profiles at the same time, a measure is desirable which reflects both the magnitude and shape similarity of the gene expression data. In this situation, the integration of these similarity measures can help to overcome the limitations of the existing measures. Sun et al. [24] constructed an integrated similarity measure (TMJ) by multiplying the Triangle and Jaccard similarity to serve as the basis of some similarity-based prediction models. However, the existing functional gene annotations are not utilized in this approach.

In this investigation, a novel computational integrated framework, called ISS, is developed by combining different similarity measures through linear combination of weights. A new fitness function, called FFFAG, is also introduced to estimate the value of ISS. In ISS, biological knowledge is incorporated in the computation of weights for each similarity measure. The novelty of this study lies in

i. developing ISS for gene expression data by combing different similarity measures for improving expression similarity
ii. a fitness function using functional annotations of genes. In the first case, the concept of integrating different similarity measures for improving expression similarity is introduced for the first time. Moreover, it is highly likely that this concept is unique for other fields also. In the second case, the fitness function computed by considering the difference of functional similarity and ISS is a new concept. The application of ISS for determining the value of weights in a similarity score (like ISS) is shown for the first time.

## 2. Background

### 2.1. Different similarity measures

The methods to calculate the similarity between the gene pairs are discussed here. Let *𝒳*= *x*_1_, *x*_2_, · · ·, *x*_*N*_ and *𝒴*= *y*_1_, *y*_2_, · · ·, *y*_*N*_ be the expression levels of two genes in terms of log-transformed microarray gene expression data where *N* is the total number of expression values.

#### 2.1.1. Euclidean distance (ED)

The ED between genes *𝒳* and *𝒴* is calculated as

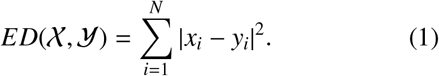

The values obtained from ED are scaled within 0 to 1 and then inverted to find the gene similarity.

#### 2.1.2. Pearson correlation (PC)

The PC between gene *𝒳* and *𝒴* is computed as

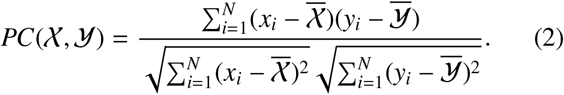

where 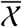 and 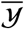 are the mean expression value of gene *𝒳* and *𝒴* gene, respectively. If *𝒳* and *𝒴* are positively correlated then *PC*(*𝒳, 𝒴*) is 1, if *𝒳* and *𝒴* are inversely correlated then *PC*( *𝒳, 𝒴*) is -1, and if *𝒳* and *𝒴* are totally uncorrelated then *PC*(*𝒳, 𝒴*) is 0. The values obtained from PC is scaled within 0 to 1 such that 0 means no correlation and 1 means highly correlated.

#### 2.1.3. Spearman rank correlation (SRC)

The SRC coefficient is the PC coefficient between the ranked variables and it is computed as

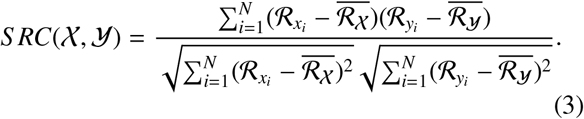

where numerator is the covariance of the ranked variables and denominator is the multiplication of standard deviations of the ranked variables. The values obtained from SRC is scaled within 0 to 1.

#### 2.1.4. Manhattan distance (MD)

The MD is defined as:

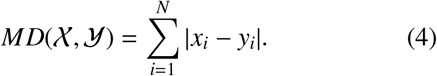

The values obtained from MD are scaled within 0 to 1 and then inverted for finding the gene similarity.

### 2.2. Computing the Similarities in a Single Framework

To precisely calculate the importance of each similarity measure by a single criterion followed by the integration of the similarity measures require an unified scoring framework. In this regard, the similarities computed from different similarity measures are re-scored individually using Saccharomyces Genome Database (SGD) annotations [25]. Genes/proteins that are related to the same process are presumed to be functionally linked. The true positive (TP) number of gene pairs at different similarity values are calculated for each similarity measure. The TP gene pairs have same Gene Ontology (GO) term. Positive predictive value (PPV) can be computed as [7]

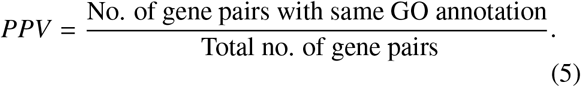

Here, the gene pairs having common gene annotation share same yeast GO-Slim process annotation and the available number of gene pairs at a specific similarity value is the total number of gene pairs. “The hierarchical nature of GO and multiple inheritance in the GO structure can lead to evaluation problems if we consider only the particular GO term with which a gene is annotated” [7]. Further, gene location and molecular activity are also included in GO terms. To simplify this problem, three different annotation procedures (biological process, location and molecular activity) are designed in Saccharomyces Genome Database [25] where every gene is annotated at the same level without any tree-based structure. In this investigation, the Yeast GO-Slim process (biological) annotations (https://www.yeastgenome.org/goSlimMapper) are considered because our ultimate aim is to predict the gene functions, not their location or molecular function.

## 3. Proposed Methodology

The key steps of our proposed methodology can be summarized as follow:

a. Calculate pairwise similarity of genes from expressions using various similarity measures.
b. Rescore the similarities in a same framework of PPV [26] using Saccharomyces Genome Database (SGD) annotations.
c. Integrate the various similarity measures in a weighted combination style where the weight for each measure is determined systematically using gene annotation.

The similarities among the genes using gene expression data, from different similarity measures, are individually calculated in the framework of PPV, based on SGD annotations [25].

### 3.0.1. Proposed Framework for Integration of Similarity Measures

The goal of this investigation is to integrate different similarity measures through a score by finding their appropriate weight using functional annotations. This integrated score should help in finding functionally similar genes which have similar expression profiles in a better fashion. An integrated score should reflect the functional similarity of the genes more appropriately compared to the existing similarity measures using their expression profiles. Some weights (*w*_1_, · · ·, *w*_*m*_) are first assigned to integrate the PPVs of different similarity measures (*𝒮*_1_, · · ·, *𝒮*_*m*_). The ISS, between two gene expressions *𝒳* = *x*_1_, · · ·, *x*_*n*_ and *𝒴* = *y*_1_, · · ·, *y*_*n*_ is defined as

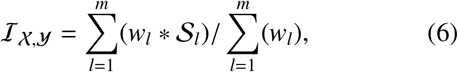

where *m* is the number of similarity measures.

The PPVs computed between two genes *𝒳* and *𝒴* using MD, ED, PC, and SRC are represented by *𝒮*_1_, *𝒮*_2_, *𝒮*_3_ and *𝒮*_4_, respectively. These PPVs are integrated through weights *w*_1_, *w*_2_, *w*_3_, and *w*_4_ in a linear combination style. The optimal weight combination for various similarity measures is determined by minimizing the fitness function (described in Section 3.0.2) computed by considering the difference between the functional similarity (ℳ ) and the ISS ( ℐ). For example, the ISS for four similarity measures can be represented as

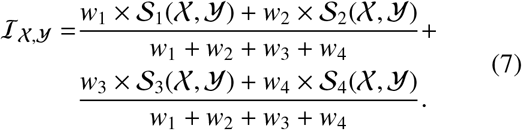

Here, the weights *w*_1_, *w*_2_, *w*_3_, and *w*_4_ vary within the range of 0.10 to 1 to find the weight combination for which the difference between ℳ and ℐ is minimum. It can be shown that:

a. 0 ≤ ℐ_*X,Y*_ ≤ 1, and
b. ℐ_*X,Y*_ = ℐ_*X,Y*_ (symmetric).

For example, two genes *𝒳* and are considered with expression values 0.21, 0.89, 0.82, 0.23 and 0.30, 0.52, 0.10, 0.90, respectively. The PPVs & optimal weight combination are 0.70, 0.10, 0.30, and 0.50 & 0.80, 0.20, 0.23, and 0.40 for MD, ED, PC, and SRC, respectively. Then the integrated score will be [(0.70 × 0.80) + (0.10 × 0.20)+ (0.30 × 0.23) + (0.50 × 0.40) / (0.80 + 0.20 + 0.23 + 0.40)] = 0.87.

### 3.0.2. Proposed Fitness Function

The grouping of genes into clusters based on expression similarity is the key established methodology to predict the function of unclassified genes [27; 9]. Therefore, if the functional similarity between two genes is high, the expression similarity between these genes should be more and vice versa. In this regard, a new fitness function, called FFFAG, is proposed to calculate the appropriate weight for each similarity measure. The fitness function is considered as the absolute difference between the functional similarity and the ISS and is defined as

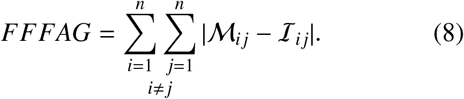

Here, *n* is the no. of genes and ℳ_*i, j*_ denotes the functional similarity between gene *g*_*i*_ and *g*_*j*_. The functional similarity value (ℳ _*i j*_) of a gene pair is 1 if both genes belong to one or more common Yeast GO-Slim categories, otherwise it becomes 0. Here, those gene pairs are only considered where both the genes in a pair are classified (i.e., both genes belong to at least one GO category).

Our goal is to minimize the fitness function (FFFAG) by varying the weights of similarity measures. The minimization of FFFAG will increase the expression similarity value of those gene pairs having higher functional similarity and will decrease the expression similarity value of those gene pairs having lower functional similarity. For a particular gene pair, the functional similarity ( ℳ_*i j*_) consists of value 0 (no functional similarity) or 1 (highly similar). If the functional similarity between the two genes is 1 and their expression similarity is 0.70, then it will contribute 0.3 (absolute difference between 1 and 0.70) to the FFFAG. In this case, an increase in expression similarity will take place through the variation of weights in ISS for those gene pairs whose functional similarity is 1. Similarly, if the functional similarity between the two genes is 0 and their expression similarity is 0.70, then it will contribute 0.70 (absolute difference between 0 and 0.70) to the FFFAG. Here, a decrease in expression similarity will take place through variation of weights in ISS for those gene pairs whose functional similarity is 0. So, the weights will be varied in such a way that the fitness function (FFFAG) will minimize. As the functional similarity (ℳ _*i j*_) for a gene pair is fixed, the increase in ISS (ℐ _*i j*_) will minimize FFFAG.

### 3.0.3. Weight Estimation in ISS using FFFAG

The weights for various similarity measures in ISS are calculated by minimizing FFFAG (difference between functional similarity using SGD annotations and ISS) through varying the weights (value of *w*_1_, *w*_2_, *w*_3_, and *w*_4_) in Eq. 7. The steps to determine the optimal weight value for each similarity measure are described below:

*S* 1) Initialize all the weight as 0.10.

*S* 2) Compute the ISS for all the gene pairs using Eq. 7.

*S* 3) Calculate the difference between functional similarity and ISS for all the gene pairs using the proposed FFFAG (Eq. 8).

*S* 4) Vary the weights in steps of 0.1 and repeat steps *S* 2 and *S* 3 to get a weight combination for similarity measures for which FFFAG is minimized.

Fig. 1 demonstrates the variation of FFFAG for different weights of All Yeast data using ISS. The estimated weight combinations of similarity measures for various datasets are as follows:

**Figure 1:**
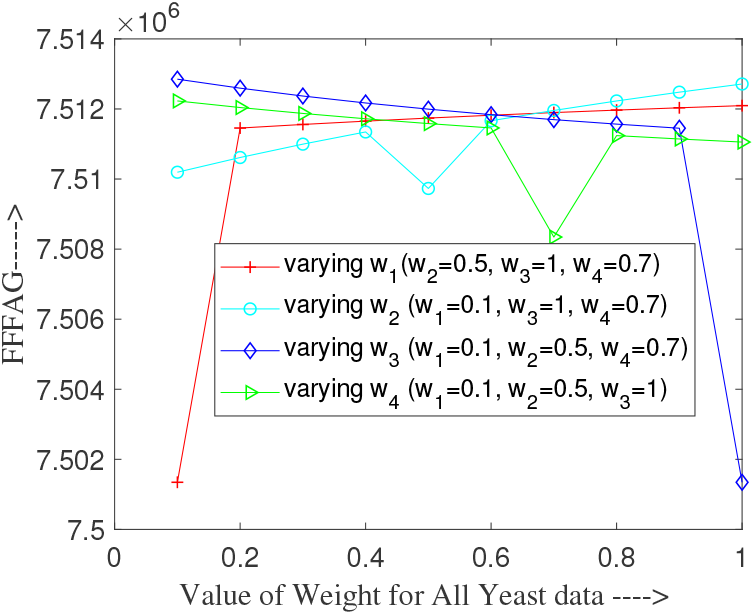
Variation of FFFAG with different weights for All Yeast data using ISS. One weight is changed whereas the other weights are fixed.

All Yeast: *w*_1_ = 0.10; *w*_2_ = 0.50; *w*_3_ = 1.0; *w*_4_ = 0.70, DSAY: *w*_1_ = 0.10; *w*_2_ = 0.10; *w*_3_ = 1.0; *w*_4_ = 0.10,

CCAY: *w*_1_ = 0.20; *w*_2_ = 0.01; *w*_3_ = 0.86; *w*_4_ = 0.24,

SAY: *w*_1_ = 0.40; *w*_2_ = 0.10; *w*_3_ = 0.80; *w*_4_ = 0.70,

Cell Cycle: *w*_1_ = 0.20; *w*_2_ = 0.01; *w*_3_ = 0.85; *w*_4_ = 0.60,

Yeast Complex: *w*_1_ = 0.50; *w*_2_ = 0.40; *w*_3_ = 0.80; *w*_4_ = 1.0.

### 3.0.4. Proposed TMJ

As mentioned earlier, Sun et al. [24] constructed an integrated similarity measure (TMJ) by multiplying the Triangle and Jaccard similarity. Functional annotation is not used for this integration. We modified the TMJ measure. Here, the Triangle and Jaccard similarities are integrated using functional annotations. The steps of the modifying TMJ are similar to the steps of ISS. The PPVs computed between two genes *𝒳* and *𝒴* using Triangle and Jaccard similarity are represented by *𝒮*_*T*_, and *𝒮*_*J*_, respectively. These PPVs are integrated through weights *w*_*T*_, and *w*_*J*_ in a linear combination style. The proposed FFFAG is used for estimating appropriate weights for Triangle and Jaccard similarity. FFFAG is minimized and it is computed by taking the difference between TMJ and functional similarity. The modified TMJ (MTMJ) can be represented as

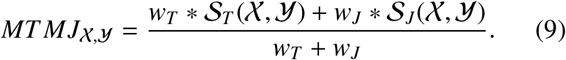

## 4. Experimental Results

The experimental results of ISS and its scope of unclassified gene’s function prediction are described in this section.

### 4.1. Dataset Description

Six *Saccharomyces cerevisiae* datasets, namely All Yeast [1], Diauxic Shift All Yeast (DSAY) [1], Cell Cycle All Yeast (CCAY) [1], Sporulation All Yeast (SAY) [1], Cell Cycle [28], and Yeast Complex [29] are employed in the present study. The name of the datasets, the no. of genes, and no. of. time points of the datasets are presented in Table 1. The missing gene expression values are predicted by LSimpute method [30] for all datasets.

**Table 1:** Details of datasets.

| Dataset name | All Yeast | DSAY | CCAY | SAY | Cell Cycle | Yeast Complex |
| --- | --- | --- | --- | --- | --- | --- |
| No. of genes | 6072 | 6072 | 6072 | 6072 | 634 | 979 |
| No. of expression values | 80 | 7 | 60 | 13 | 184 | 79 |

### 4.2. Performance Evaluation

The performance of ISS is assessed based on the following:

i. Plotting PPVs vs. similarity values (Fig. 2a-2d), and
ii. Plotting PPV vs. top gene pairs (Fig. 3a-3a).

**Figure 2:**
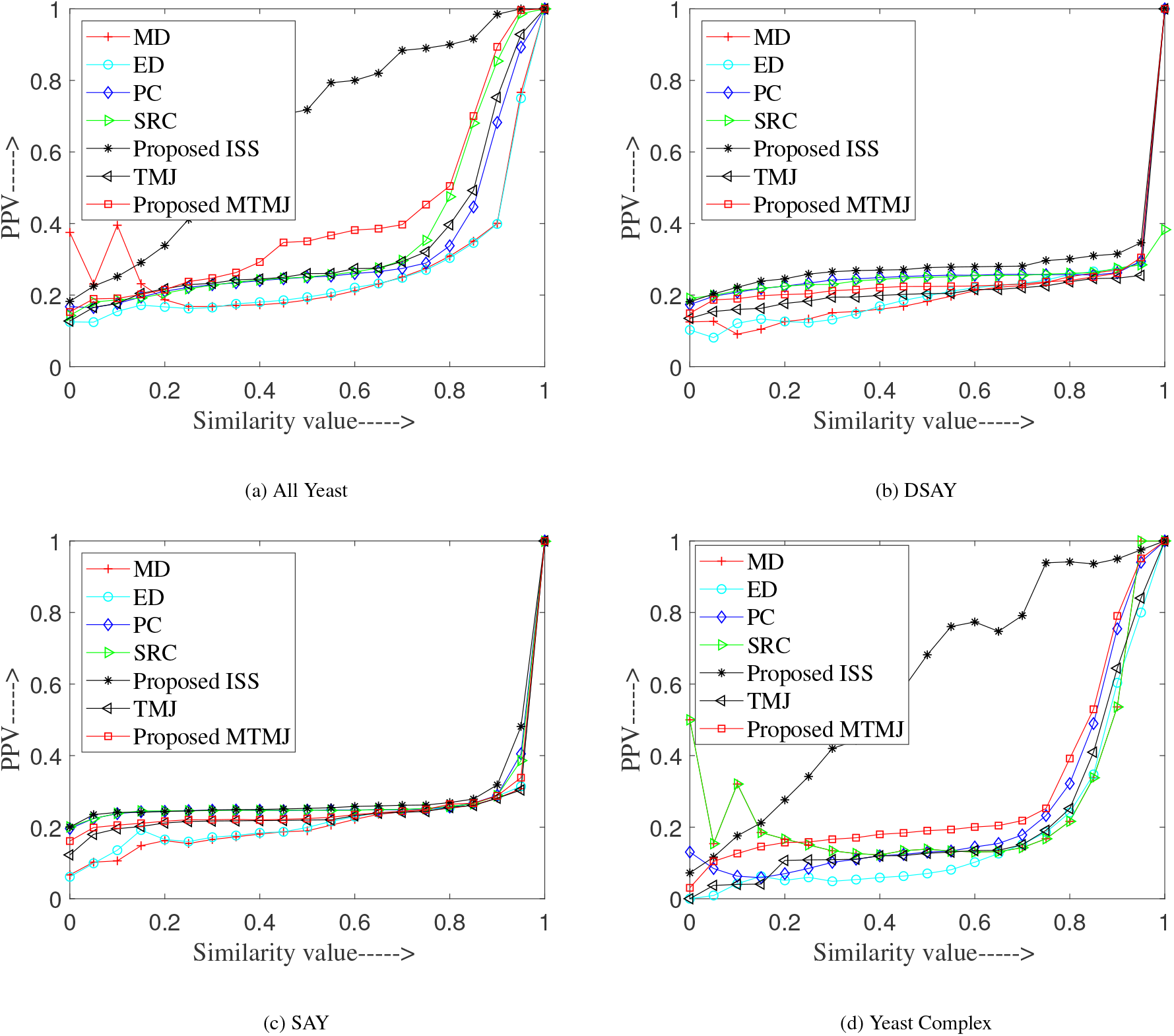
Variation of PPVs across similarity values for ISS and various similarity measures.

**Figure 3:**
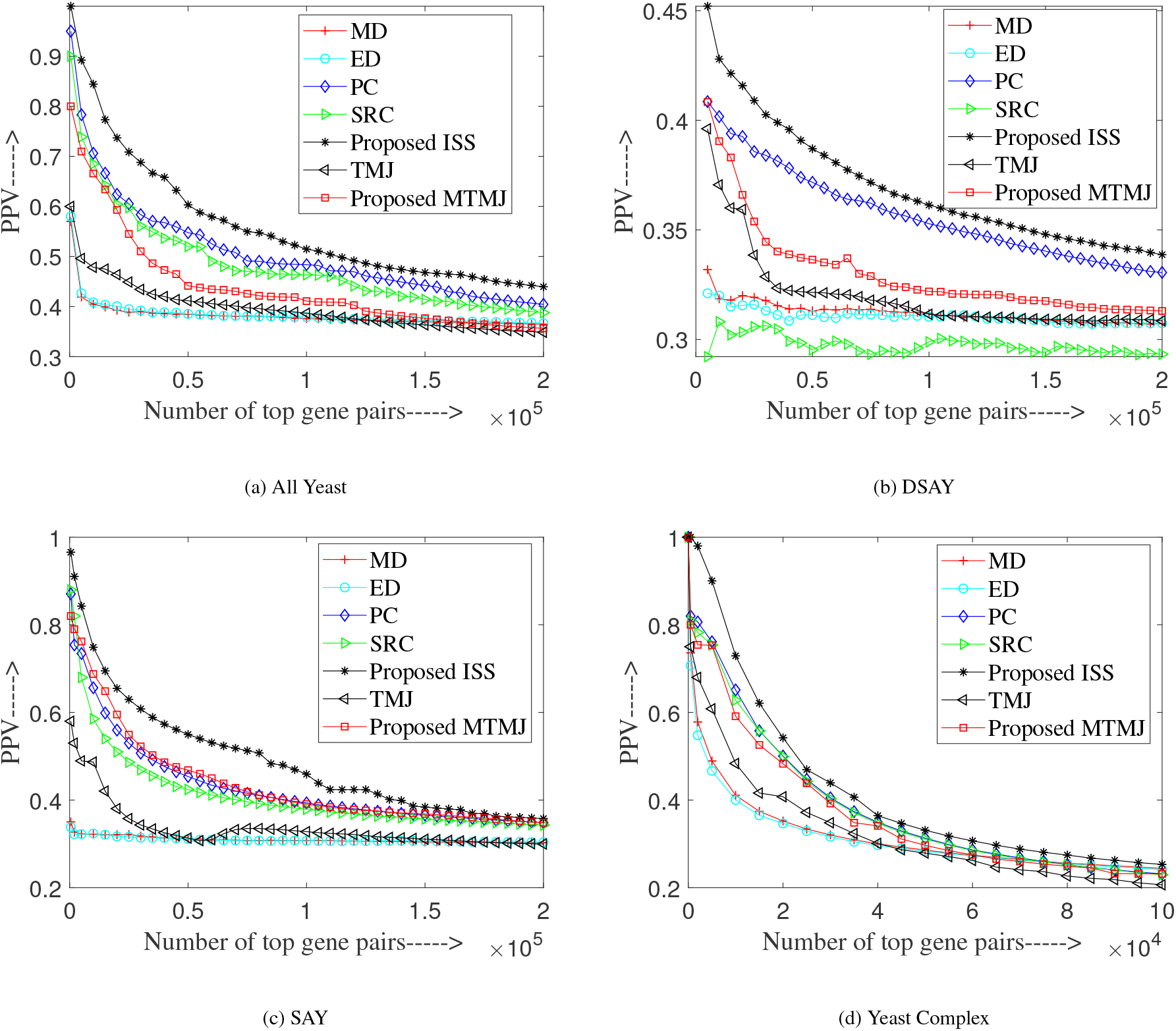
variation of PPVs across top gene pairs between ISS and different similarity measures.

#### 4.2.1. PPV and similarity values

The PPV values for the various similarity values are calculated for the proposed method ISS and the existing measures using All Yeast, DSAY, SAY, Yeast Complex, CCAY, and Cell Cycle datasets. The figures for the first 4 datasets are presented in Fig. 2a-2d, whereas the same for the last two datasets are available in the supplementary file http://www.isical.ac.in/∼shubhra/supplimentaryFile_new.pdf. It is evident that the curves attributed to ISS appear at the top as compared to the existing measures as the expression similarity between gene pairs are improved by integrating different similarity measures. For example, at 0.85 similarity value, the PPV values of the MD, ED, PC, SRC, TMJ, MTMJ, and ISS are 0.35, 0.34, 0.44, 0.68, 0.49, 0.70, and 0.92, respectively for All Yeast dataset. Notably, the curves corresponding to MD and ED always lie at the bottom (performance are inferior). We observed that the proposed MTMJ’s performance is improved from the existing TMJ measure.

#### 4.2.2. PPV and top gene pairs

Performance of ISS in case of PPV vs. top gene pairs for all the datasets is compared with existing measures. The curve for the same datasets are presented in Figs. 3a-3d. For CCAY and Cell Cycle datasets the figures are presented in the supplementary file. We noticed that the relationship between PPV and top gene pairs no. is inversely proportional. Further, for all the datasets, the result for the proposed ISS is noticeably superior as the curve corresponding to ISS always lies at the top. For example, the PPV values for MD, ED, PC, SRC, TMJ, MTMJ, and ISS are 0.38, 0.38, 0.54, 0.52, 0.41, 0.44, and 0.61, respectively for All Yeast dataset using top 50,000 gene pairs. Notably, the curves corresponding to MD and ED lie at the bottom. The proposed MTMJ’s performance is noticeably enhanced than the existing TMJ.

### 4.3. Cross-Validation

In this investigation, 5-fold cross-validation is employed to portray the performance of ISS in a conventional way. Here, 4 folds out of 5 folds of the genes are used for training and the remaining one fold is used for the performance evaluation. In the first (training) phase, the weight values are determined for different similarity measures by minimizing the difference between the functional similarity and the ISS. In the second (evaluation) phases, the weights are assigned to the similarity values of different similarity measures obtained from the expressions of the remaining one fold. The performance of ISS, which was obtained using test set is compared with the existing measures. For each dataset, this is repeated 5 times and the PPV vs. similarity value results for one of those processes are presented in Figs. 4a-4d. The cross-validation result for variation of PPV across similarity value for CCAY and Cell Cycle datasets and variation of PPV across top gene pairs for all the datasets are available in the supplementary file. It is clear from the figures that the ISS’s performance is better than existing measures as the curves for ISS lies above the curves.

**Figure 4:**
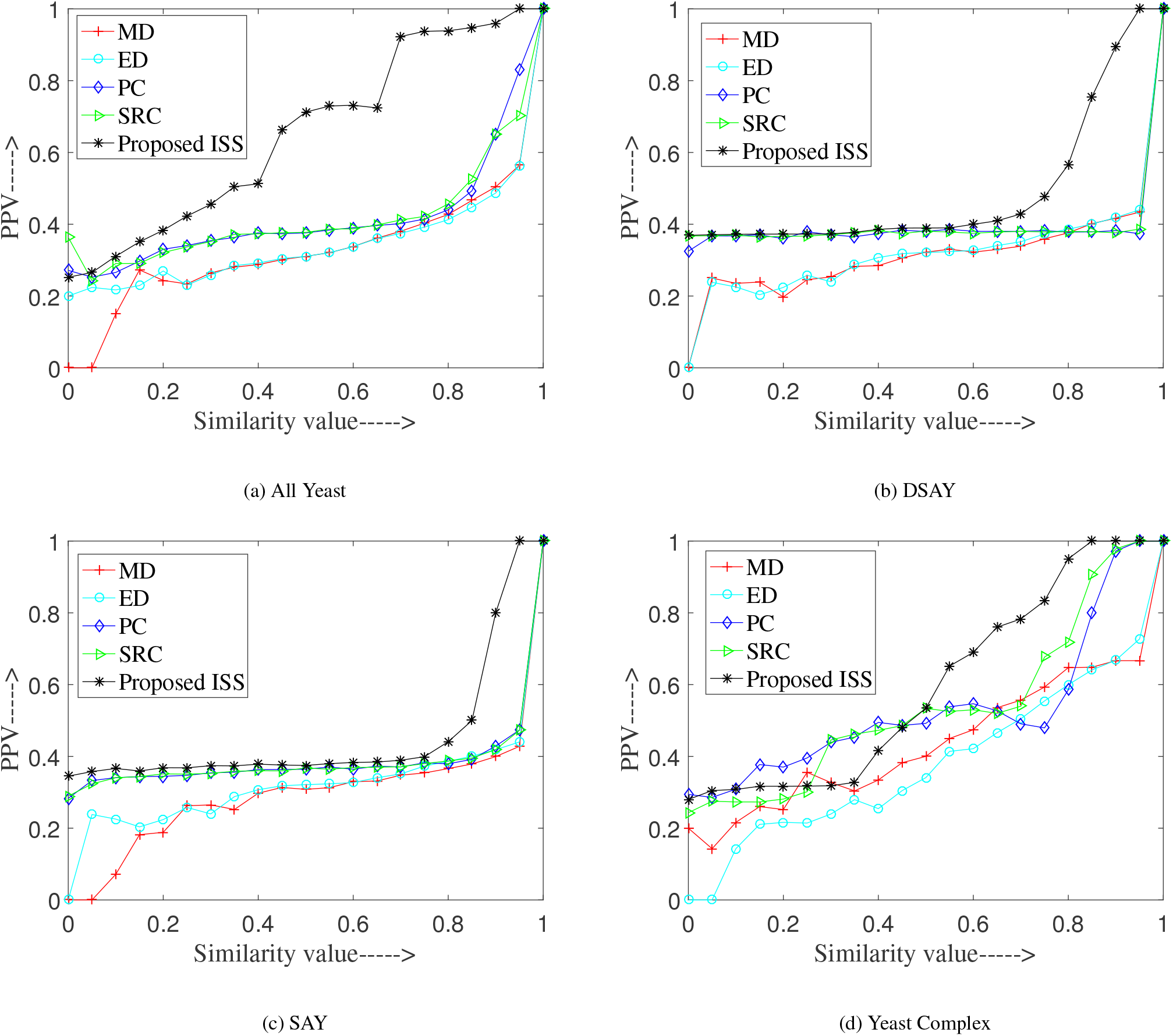
Cross-validation results comparing ISS with existing measures in PPVs and similarity values.

### 4.4. Function Prediction

All Yeast dataset and MIPS annotation [31] are employed to classify unclassified genes. Here, *k*-medoids [32] is used to cluster the similarity score achieved by the ISS.

From the functional enhancement of the cluster, the biological function of an unknown gene is predicted The steps of function prediction of unknown genes are as follows:

*S* 1) Compute the ISS for all possible pairs by assigning weight to each similarity measure using proposed method.

*S* 2) Group the genes by *k*-medoids clustering.

*S* 3) Identify the clusters with functional enrichment in MIPS categories. In this step, 12 clusters are found with *p*-value cutoff 10^−10^.

*S* 4) Predict functions of unclassified genes from functionally enriched clusters. Here, function of 40 unclassified yeast genes are predicted.

Table 2 presents function predictions of 40 unclassified genes with a *p*-value cutoff of 10^−10^ using MIPS annotations for the All Yeast data.

**Table 2:** Predicted function of 40 unclassified genes from 12 clusters using MIPS annotations and All Yeast dataset at *p*-value cutoff 10^−10^.

| Cluster number | Unclassified genes | Functional category | $p$-value | No. of genes within cluster | No. of genes within the genome |
| --- | --- | --- | --- | --- | --- |
| 1 | YDR493W YHR059W<br>YLR204W YML030W<br>YLR253W | mitochondrion | 1.4238e-16 | 17 | 154 |
| 2 | YGR210C | ribosomal proteins | 3.6283e-15 | 16 | 221 |
| 3 | YCR016W YLR287C<br>YDR161W YIL110W<br>YDR198C | rRNA processing | 5.4685e-14 | 16 | 167 |
| 4 | YER158C | electron transport and membrane-associated energy conservation | 2.8368e-16 | 13 | 48 |
| 5 | YGR154C YDR374C<br>YOL131W YLR445W<br>YLR376C YGL081W<br>YGR053C YHR079BC<br>YFL051C | meiosis | 2.1426e-16 | 17 | 156 |
| 6 | YIR016W YBR151W | protein/peptide degradation | 2.6172e-18 | 18 | 251 |
| 7 | YOR007C YKR018C<br>YOL057W YJL021C<br>YOR258W | protein folding and stabilization | 1.4889e-15 | 12 | 91 |
| 8 | YPR202W YPR203W<br>YBL110C YLR464W | DNA topology | 3.2404e-14 | 9 | 52 |
| 9 | YER0191C | protein synthesis | 8.5075e-17 | 30 | 453 |
| 10 | YCL056C | proteasomal degradation (ubiquitin/proteasomal pathway) | 2.4129e-15 | 11 | 126 |
| 11 | YOR164C YOL098C<br>YCLX05C | proteasomal degradation (ubiquitin/proteasomal pathway) | 7.4212e-11 | 9 | 126 |
| 12 | YLR196W YLR051C<br>YOR051C | rRNA processing | 1.5667e-11 | 11 | 167 |

### 4.5. Biological Importance

In this section, the biological importance of function prediction for the unknown genes will be discussed (Table 2).

The function for gene YLR204W in cluster 1 is predicted as “mitochondrion”. The mitochondrial gene COX1 is generally associated with multiple exons and introns in most strains of *Saccharomyces cerevisiae*. The processing of COX1 primary transcript requires accessory proteins factors out of which some are encoded by nuclear genes and others by reading frames residing in some of the introns of the COX1 and COB genes. It was shown that the gene YLR204W encodes a protein which is involved in the processing of COX1 RNA intermediate [33]. Therefore, predicting the function of gene YLR204W as “mitochondrion” is a promising one.

The function of gene YDR374C in cluster 6 is predicted as ‘meiosis”. The gene YDR374C falls among the 82 core Ume-6 regulated genes, out of which 52 genes encode proteins with known function. Surprisingly, half of them (26 out of 52 genes) are known to play key roles in meiosis [34]. In addition, YDR374C recently reported as needed for effective sporulation [34]. Hence, the function prediction for the gene YDR374C as “meiosis” is a possible one.

The predicted function for YOR258W in cluster 7 is “protein folding and stabilization”. The gene YOR258W encodes a protein which is a homologue to Aprataxin, a protein that is encoded by the APTX gene. In [35], it is reported that APTX impacts protein folding. Therefore, predicting the function of gene YOR258W as “protein folding and stabilization” is a likely one.

## 5. Discussion

A method for integrating different similarity measures, named ISS, is developed using gene annotations for truly reflecting their expression similarity in accordance with their functional similarity. The current study employed six *Saccharomyces cerevisiae* datasets. We observed that integrated score (ISS) is noticeably enhanced for PPVs at different similarity values (presented in Figs. 2a-2d) as well as for several top gene pairs (presented in Figs. 3a-3d) as compared to those of the existing individual similarity measures such as PC [16; 36], MD [3], ED [17], and SRC [18]. We have seen from the figures that the curves for proposed ISS appear above the other curves for both types of criteria (PPV vs. similarity measures and PPV vs. top gene pairs). Hence, it can be concluded that the integration of existing similarity measures using functional annotations improved expression similarities as compared to those of individual measures for all the datasets.

In addition, the predicted functions of unclassified genes are validated with existing biological literatures. For example, the predicted function of gene YLR204W as “mitochondrion” appears to be likely because it encodes a protein that is involved in the processing of the mitochondrial gene COX1 RNA intermediate. The gene YDR374C is validated to be involved in “meiosis”. The predicted function for YOR258W as “protein folding and stabilization” is also validated.

## 6. Conclusion

In summary, ISS is developed by integrating various similarity measures for improving expression similarity of genes based on functional annotations. First, the pairwise expression similarity for all possible gene pairs is computed using the existing similarity measures. Expression similarity for different measures is then converted to a single framework of PPV, involving functional annotations. The PPV of each measure is integrated through a similarity score by assigning an appropriate weight to each measure. A new fitness function, called FFFAG, is also developed by minimizing the difference between functional similarity value and the ISS. The new FFFAG is used to determine the linear combination of weights for different similarity measures in ISS. Here, the functional similarity is measured using functional annotations available for the genes. An existing similarity measure, TMJ, is also modified to incorporate biological knowledge involving functional annotation and it is compared with ISS.As SGD annotations for genes are employed to estimate weight combinations, MIPS annotations are applied for evaluation purposes. Functions of 40 unclassified yeast genes are then predicted from 12 clusters at *p*-value less than 10^−10^ by using the *k*-medoids clustering technique on the ISS. An improved performance of the developed method is evident from identifying the similarity between the gene expression profiles in terms of functional annotations.

The ISS is appeared to be helpful in functions prediction of unclassified gene using *k*-medoids clustering. Here, the function is predicted using MIPS annotations, where a gene exists in one cluster. It may be a new research direction if fuzzy clustering method is used instead of *k*-medoids, where a gene can exists in many clusters instead of single cluster with fuzzy membership.

## Declarations

### Data Availability

Data will be made available on reasonable request from first author.

